# Experimental introgression in *Drosophila*: asymmetric postzygotic isolation associated with chromosomal inversions and an incompatibility locus on the X chromosome

**DOI:** 10.1101/2022.07.07.499141

**Authors:** N. Poikela, D. R. Laetsch, M. Kankare, A. Hoikkala, K. Lohse

## Abstract

Interspecific gene flow (introgression) is an important source of new genetic variation, but selection against it can reinforce reproductive barriers between interbreeding species. We used an experimental approach to trace the role of chromosomal inversions and incompatibility genes in preventing introgression between two partly sympatric *Drosophila virilis* group species, *D. flavomontana* and *D. montana*. We backcrossed F_1_ hybrid females from a cross between *D. flavomontana* female and *D. montana* male with the males of the parental species for two generations and sequenced pools of parental strains and their reciprocal 2^nd^ generation backcross (BC_2_mon and BC_2_fla) females. Contrasting the observed amount of introgression (mean hybrid index, HI) in BC_2_ female pools along the genome to simulations under different scenarios allowed us to identify chromosomal regions of restricted and increased introgression. We found no deviation from the HI expected under a neutral null model for any chromosome for the BC_2_mon pool, suggesting no evidence for genetic incompatibilities in backcrosses towards *D. montana*. In contrast, the BC_2_fla pool showed high variation in the observed HI between different chromosomes, and massive reduction of introgression on the X chromosome (large X-effect). This observation is compatible with reduced recombination combined with at least one dominant incompatibility locus residing within the X inversion(s). Overall, our study suggests that genetic incompatibilities arising within chromosomal inversions can play an important role in speciation.

## Introduction

Interspecific gene flow (introgression) is an important source of genetic variation for adaptation to new environments (Abbott et al., 2013; Anderson & Hubricht, 1938; Lewontin & Birch, 1966). At the same time, selection against introgression at certain loci acts to maintain barrier loci and protect species’ integrity from the negative effects of hybridization (Barton & Bengtsson, 1986; Ravinet et al., 2017; Servedio & Noor, 2003; Wu, 2001). The patterns of genomic divergence and the permeability of species boundaries in certain genomic regions provide valuable insights into the genomic regions that contribute to speciation (Harrison & Larson, 2014). However, we still lack a good understanding of how barrier genes are arrayed within the genome, how effectively and in what generation they restrict introgression, and what kind of role chromosomal inversions and sex chromosomes play in maintaining genetic barriers (Butlin, 2005; Coughlan & Matute, 2020; Coyne & Orr, 2004; Faria & Navarro, 2010; Gompert, Lucas, Nice, & Buerkle, 2012; Nosil & Feder, 2012).

Speciation in isolation (allopatry), occurring via drift or indirect effects of selection, can lead to the “incidental” establishment of intrinsic genetic incompatibilities (Coyne & Orr, 2004; Tang & Presgraves, 2009). These incompatibilities generally involve negative epistatic interactions between two or more loci, where new alleles arising in one or both of the interacting lineages function well in their own genetic background, but interact negatively with the alleles of other species in hybrids (Bateson-Dobzhansky-Muller incompatibilities, BDMIs or DMIs; Coyne & Orr, 2004; Orr, 1995; Presgraves, 2010b). Lack of gene flow may also increase the fixation probability of meiotic drive loci (loci that manipulate meiotic process to favour their own transmission) and their suppressors within each population and drive the genomic divergence of these populations (Crespi & Nosil, 2013). Compared to allopatric speciation, where both BDMIs and neutral differences between species are expected to build up randomly along the genome, divergence with gene flow leads to clusters of species- or population-specific loci that are sheltered from recombination (Abbott et al., 2013; Butlin, 2005; Felsenstein, 1981). Accordingly, an accumulation of BDMIs between species may be drastically different with and without gene flow. Importantly, in the presence of gene flow BDMIs can only accumulate if they are favoured by selection (Bank, Bürger, & Hermisson, 2012).

Chromosomal inversions are a major factor rearranging the genome and gene order, and inducing changes in recombination rates, gene interactions and expression patterns (Hoffmann & Rieseberg, 2008; Kirkpatrick & Barton, 2006; Sturtevant, 1921; Dobzhansky, 1940). Inversions may gain a fitness advantage and spread through conspecific populations, if they reduce recombination within co-adapted gene complexes important in adaptation and/or in maintaining species integrity (Kirkpatrick & Barton, 2006; Navarro & Barton, 2003). Once inversions have become fixed between the species, they can generate postzygotic isolation and limit gene flow between the species through problems in gamete formation and/or in the build-up of BDMIs. Single recombination events (cross-overs) within paracentric inversions (breakpoints on different sides of the centromere) can produce malformed gametes with dicentric and acentric chromosomes (Coyne & Orr, 2004; Hoffmann & Rieseberg, 2008; Rieseberg, 2001). However, in *Drosophila* the problems with malformed gametes are partially avoided, since these gametes remain in the polar nuclei and do not enter the developing gametes (Hoffmann & Rieseberg, 2008; Sturtevant & Beadle, 1936). Perhaps more importantly, reduced recombination across inverted regions, particularly near inversion breakpoints and within overlapping inversions, facilitates the build-up of BDMIs via divergent selection and/or drift (Fishman, Stathos, Beardsley, Williams, & Hill, 2013; Khadem, Camacho, & Nóbrega, 2011; Mcgaugh & Noor, 2012; Navarro & Barton, 2003; Noor, Grams, Bertucci, & Reiland, 2001). While blocks of genetic material can occasionally be exchanged through double cross-overs within long inversions (Navarro, Betrán, Barbadilla, & Ruiz, 1997) and smaller DNA sections (several hundred bps) though gene conversion events within any kind of inversions (Korunes & Noor, 2018), recombination within inversions generally remains lower than on colinear chromosome sections (Hoffmann & Rieseberg, 2008). Thus, species-specific inversions harbouring BDMIs may act as strong barriers to gene flow (Hoffmann & Rieseberg, 2008; Noor et al., 2001).

The disproportionate involvement of sex chromosomes in reproductive isolation in many systems is captured by two general observations: Haldane’s rule – the increased F_1_ inviability and sterility of the heterogametic sex compared to the homogametic sex (Haldane, 1922; Orr, 1997; Turelli & Orr, 2000) – and the large X-effect – the fact that the X chromosome shows a disproportionately large effect on the sterility and inviability of backcross hybrids (Masly & Presgraves, 2007; Turelli & Orr, 2000). Explanation for both observations often presume recessivity of X-linked alleles, which can lead to more pronounced effects in hemizygous than in heterozygous hybrids (“Dominance theory”; Coyne & Orr, 2004; Turelli & Orr, 1995, 2000) and/or rapid evolution of X-linked alleles facilitating BDMIs as a byproduct (“Faster X evolution”; Charlesworth, Campos, & Jackson, 2018; Charlesworth, Coyne, & Barton, 1987). The X chromosome has also been suggested to be enriched for genes that create postzygotic isolation in hybrids compared to autosomes (Coyne, 2018). In particular, meiotic drive loci are more frequent on the X than on autosomes, and incompatibilities between drivers and their suppressors in hybrids may generate problems in hybrid development (Courret, Chang, Wei, Montchamp-Moreau, & Larracuente, 2019; Crespi & Nosil, 2013; Crown, Miller, Sekelsky, & Hawley, 2018).

Pairwise BDMIs may involve substitutions in both diverging lineages, or derived substitutions in one lineage and preserved ancestral alleles in another lineage (Barbash, Awadalla, & Tarone, 2004; Cattani & Presgraves, 2009; Coyne & Orr, 2004). BDMIs can also result from cumulative effects of many small incompatibilities or from a single incompatibility between two complementary genes, and the complexity of the incompatibility interaction does not reflect the severity of the barrier (Orr, 1995; Presgraves, 2010a). Importantly, and in contrast to interactions within a locus where a dominant allele masks a recessive allele, in epistatic interactions between different loci a dominant allele at one locus may interact with dominant or recessive alleles at other loci. Epistatic interactions involving dominant alleles are of special interest in the context of BDMIs, but they have received less attention than BDMIs involving recessive alleles.

Two closely-related species of the *Drosophila virilis* group, *D. montana* and *D. flavomontana*, provide an excellent test case for studying the evolution of BDMIs. The species originate from the Rocky Mountains of North America, where the divergence of the *montana* complex species (*D. flavomontana, D. lacicola* and *D. borealis*) most likely occurred (Hoikkala & Poikela, 2022; Patterson, 1952; Throckmorton, 1982). *D. montana* has expanded around the northern hemisphere, whereas *D. flavomontana* has remained in North America (Hoikkala & Poikela, 2022). *D. montana* lives generally in colder environments and uses different host trees than *D. flavomontana* (Patterson, 1952; Throckmorton, 1982). Reproductive barriers between *D. montana* females and *D. flavomontana* males are nearly complete, with extremely strong prezygotic barriers and inviability and sterility of rarely produced F_1_ hybrids (Noora Poikela et al., 2019). However, in crosses between *D. flavomontana* females and *D. montana* males, strong postzygotic isolation is accompanied by prezygotic barriers of variable strength, and F_1_ hybrid females can still be crossed with the males of both parental species to obtain backcross progenies in both directions (Noora Poikela et al., 2019). Interspecific hybrids have also reportedly been found in nature (Patterson, 1952; Throckmorton, 1982). Our recent demographic modelling shows that the species have diverged ~3 Mya, with low levels of postdivergence gene flow from *D. montana* to *D. flavomontana* (Poikela, Laetsch, Lohse, & Kankare, 2022). Moreover, we found several inversions that were fixed between the species in all studied individuals across different populations in North America (Poikela et al., 2022). These inversions were already present in species’ common ancestor, and they may have contributed to the build-up and maintenance of adaptive traits and reproductive barriers by restricting gene flow between the evolving lineages (Poikela et al., 2022).

The goal of this study was to determine which genomic regions are likely to accommodate dominant BDMIs in hybrids between *D. montana* and *D. flavomontana*, paying special attention to fixed inversions and the X chromosome. We investigated BDMIs between these species experimentally by sequencing pools of *D. montana* females from an allopatric population and *D. flavomontana* females from a (presently) parapatric population, as well as pools of 2^nd^ backcross generation (BC_2_) females in both directions (Fig. 1). We identified chromosomal regions with decreased and increased introgression by quantifying the amount of introgressed genetic material (mean hybrid index, HI) along the genome in both backcross pools. We then compared the observed HI to the distribution of chromosome-wide HI in *in silico* replicates of this “introgress-and-resequence” experiment under contrasting assumptions about the presence and location of BDMIs. Since this experimental design involved backcross females, we were able to detect only BDMIs involving a dominant allele, while the recessive-recessive BDMIs remained masked (Table 1). Our main questions were:

i. Does the strength and genomic distribution of genetic incompatibilities between *D. montana* and *D. flavomontana* differ between the reciprocal crosses?
ii. Do the species show increased genetic divergence and decreased introgression within chromosomal inversions, and could this be caused by inversions’ propensity to suppress recombination and harbour genetic incompatibilities?
iii. Does the X chromosome show less introgression than autosomes (large X-effect)? And if yes, why?

**Figure 1.**
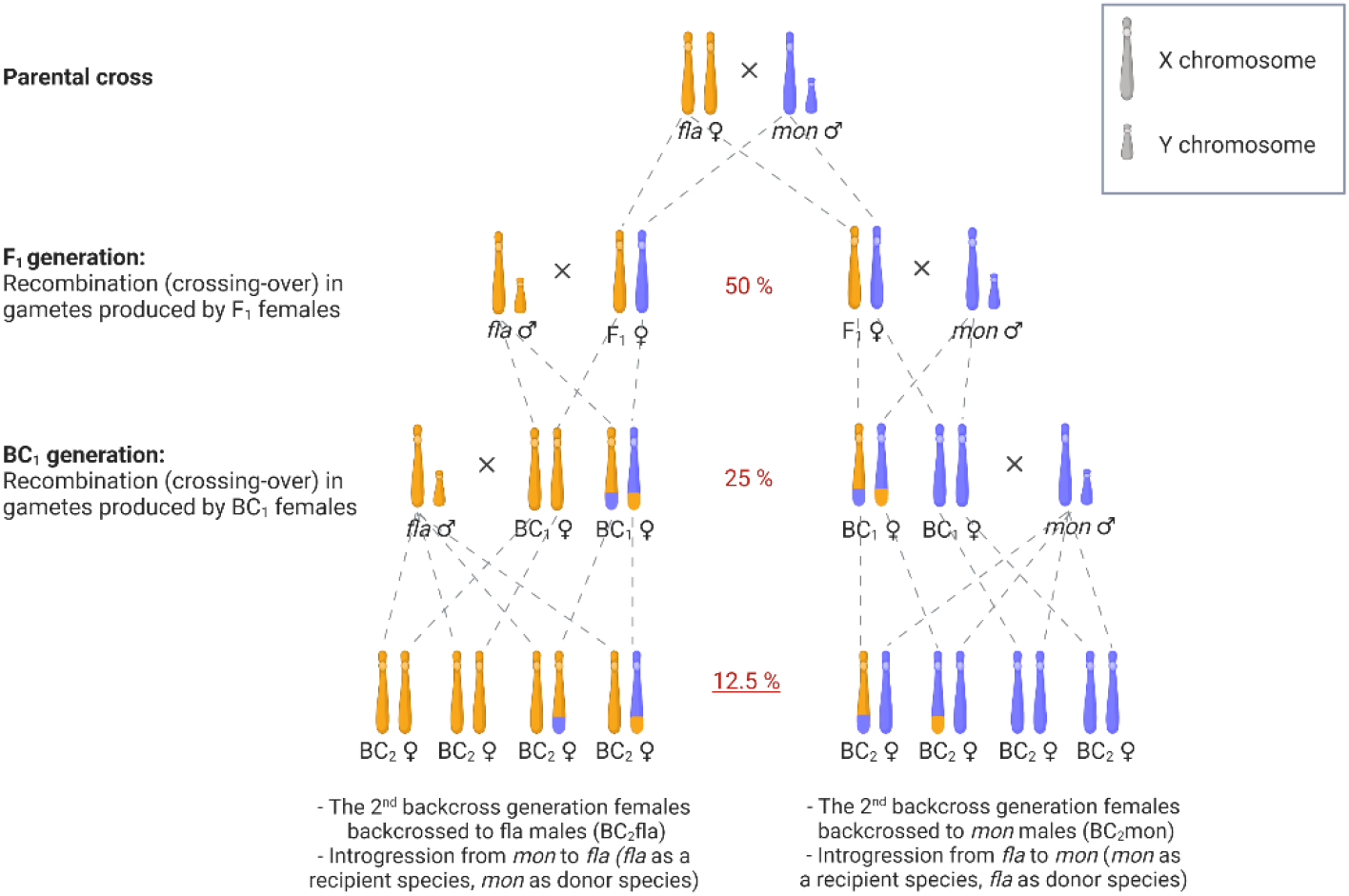
Illustration of the crossing experiment showing the inheritance of sex chromosomes (inheritance of autosomes is similar to that of female X chromosomes). F_1_ females, produced in a single-pair cross between *D. flavomontana (fla*) female and *D. montana (mon*) male, were backcrossed to either *D. flavomontana* or *D. montana* male. In the next generation, each BC_1_ female was mated with a male of its paternal species. In every generation, the expected amount of genetic material that is transferred from the gene pool of one species into the gene pool of another one (introgression) is halved (red percentages). Thus, under a null neutral model, we expect a mean HI of 12.5 % for the BC_2_ pools that were sequenced. Note that recombination occurring in the gametes produced by F_1_ and BC_1_ females creates variation in the expected amount of HI. For simplicity, the figure shows products of only one cross-over event that has occurred in each backcross direction.

**Table 1.**
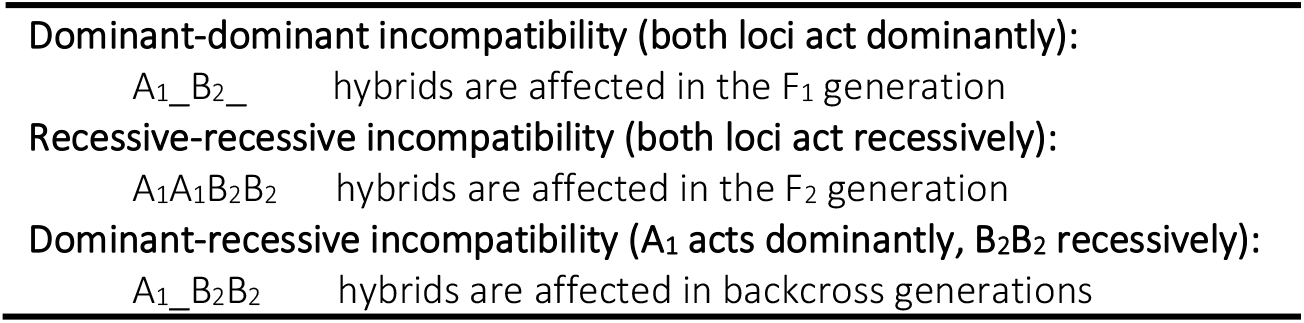
BDM model for incompatibilities (see Coyne & Orr, 2004). Here gene A1 of one species interacts negatively with gene B_2_ of another species. Underscore represents any allele, and it does not change the outcome. Note that dominance refers to an allele’s effect on fitness on a hybrid genetic background, and it does not necessarily assume dominance of alleles on their normal background within species.

## Materials and methods

### Fly material

We collected fertilised *D. montana* females from Seward, Alaska, USA (60°09’N; 149°27’W) and *D. flavomontana* females from Livingston, Montana, USA (45°20’N; 110°36’W) in 2013. The distance between the sites is ~3000 km. Alaskan *D. montana* can be regarded as an allopatric population, as *D. flavomontana* has not been found above 54°N (Noora Poikela et al., 2019). In contrast, *D. flavomontana* population from Montana can be regarded as a parapatric, as the two species are known to coexist in the Rocky Mountains, even though we found only *D. flavomontana* on the collecting site (Noora Poikela et al., 2019). We maintained the strains established from the progenies of single wild-caught *D. montana* and *D. flavomontana* females in continuous light and 19 °C for about 23 generations (~3 years) in the University of Jyväskylä (Finland) prior to their use in the present study. For the crosses, the flies were sexed under light CO_2_ anaesthesia within three days after emergence, when they were still virgins. Males and females were transferred into fresh malt-vials once a week and used in the crossing experiments at age 20 ± 2 days when they were sexually mature (Salminen & Hoikkala, 2013).

### Crossing experiment

We started the crossing experiment by performing a single-pair cross between *D. flavomontana* female (strain MT13F_1_1) and *D. montana* male (strain SE13F37), as reciprocal cross is not successful. Our crossing design (outlined in Fig. 1) only involved hybrid females because F_1_ males are largely sterile (Päällysaho, Aspi, Liimatainen, & Hoikkala, 2003; Noora Poikela et al., 2019), and because *Drosophila* males lack recombination (crossing-over) in meiosis. The initial cross produced seven F_1_ females, which were backcrossed towards both parental species: four were mated to *D. montana* males and three to *D. flavomontana* males. The 1^st^ backcross generation females (BC_1_mon and BC_1_fla females) were backcrossed to the same paternal species as in the previous generation to obtain BC_2_mon and BC_2_fla females (82 females in both directions). BC_2_ females were collected within three days after their emergence and stored in −20 °C for DNA extractions.

### Fertility of BC_1_ females

We defined the fertility of BC_1_ females by checking whether they produced progeny after mating with a *D. montana* or *D. flavomontana* male (Fig. 1). Each BC_1_ female was placed in a malt vial with a single male of either species. Once the flies mated, the couple was kept together in the vial so that the female could remate and lay eggs until she died. BC_1_ females were considered fertile, if they produced at least some larval, pupal, and/or adult-stage offspring (1=fertile, 0=sterile). We used a one-sample Student’s t-test (*t-test* function) to test whether the BC_1_ females from the reciprocal crosses showed reduced fertility, when the expected fertility was 1. We also compared the fertility of BC_1_ females between the reciprocal crosses to define possible asymmetries (BC_1_mon vs. BC_1_fla), using a generalised linear model (GLM) with Binomial distribution (1=fertile, 0=sterile) (*glm* function). All analyses were conducted in base R v1.2.1335-1 and R studio v3.6.1.

### Pool-sequencing, mapping, and variant calling

We made DNA extractions from four pools, one pool of each parental strain (*D. montana* SE13F37 and *D. flavomontana* MT13F_1_1) and pools for the two 2^nd^ generation backcrosses (BC_2_mon and BC_2_fla). Each pool consisted of 82 females. We used cetyltrimethylammonium bromide (CTAB) solution with RNAse treatment, Phenol-Chloroform-Isoamyl alcohol (25:24:1) and Chloroform-Isoamyl alcohol (24:1) washing steps and ethanol precipitation. Nextera library preparation and 150 bp Illumina paired-end sequencing were performed on two lanes using HiSeq4000 Illumina instrument at Edinburgh Genomics, UK. Illumina paired-end reads of all four samples were quality-checked with FastQC v0.11.8 (Andrews 2010) and trimmed for adapter contamination and low-quality bases using fastp v0.20.0 (using settings --detect_adapter_for_pe, --cut_front, --cut_tail, --cut_window_size 4, --cut_mean_quality 20) (Chen, Zhou, Chen, & Gu, 2018). After filtering, the total number of reads per pool varied from 153 to 174 million, the mean length and insert size peak being 141-143bp and 150bp, respectively (Table S1).

To consider potential effects of reference bias on the results, we performed the analyses using both *D. flavomontana* and *D. montana* chromosome-level reference genomes (Poikela et al., 2022). The genomes cover most regions for all the chromosomes, except for the 6th dot chromosome, and the total length of *D. flavomontana* genome is 142 Mb and that of *D. montana* 146 Mb. Filtered Illumina reads of each sample were mapped to the unmasked reference genomes using BWA mem (Burrows-Wheeler Aligner) v0.7.17 with read group information (Li & Durbin, 2009). The alignments were sorted with SAMtools v1.10 (Li et al., 2009) and PCR duplicates marked with sambamba v0.7.0 (Tarasov, Vilella, Cuppen, Nijman, & Prins, 2015). The separate BAM-files of each sample were merged and filtered for mapping quality of >20 using SAMtools. The mean coverage of the pools varied from 163 to 193 based on *D. flavomontana* reference, and 151 to 204 based on *D. montana* reference (Table S1). Allele counts for each sample at each genomic position were obtained with SAMtools mpileup using options to exclude indels and to keep reads with a mapping quality of >20 and sites with a base quality of >15. The resulting BAM-files were used for variant calling with the unmasked version of the reference genomes using heuristic SNP calling software PoolSNP (Kapun et al., 2020). In PoolSNP, we specified a minimum count of 5 to call a SNP, and a minimum coverage of 80 to reliably calculate allele frequencies and to minimize potential reference bias. For a maximum coverage, we considered positions within the 95% coverage percentile for a given sample and chromosome. Variant calling detected a total of 4,489,437 biallelic SNPs when using *D. flavomontana* reference genome, and 4,407,029 biallelic SNPs when using *D. montana* reference genome.

### Inversion breakpoints

The breakpoints of fixed inversions between *D. montana* and *D. flavomontana* on the X chromosome and chromosomes 2L, 4 and 5 were obtained from Poikela et al. (2022). The presence of the inversions in Illumina samples of parental pools was verified by passing the respective BAM-files to Delly v0.8.1 (Rausch et al., 2012), which identifies structural variants based on paired-end read orientation and split-read evidence. The inversion breakpoints were also confirmed visually by checking the orientation and insert size around each breakpoint in the Interactive Genomics Viewer (Thorvaldsdóttir, Robinson, & Mesirov, 2012) (Example plot shown in Fig. S1). Inversion breakpoints are shown in Fig. 3–4; Table S2; Fig. S3-S6).

### Genetic differentiation, hybrid index and the types of genetic incompatibilities

The expected amount of genetic material transferred from one species into the other halves with every backcross generation (Fig. 1). Given species-specific alleles, we can measure introgression via the hybrid index (HI), which can be defined simply as the heterospecific fraction of genome in an individual (or a pool of individuals). Thus, in the pool of 2^nd^ backcross generation hybrid females, the genome-wide HI is expected to be 12.5% in the absence of BDMIs (Fig. 1). However, given the random inheritance of chromatids in gametes and the randomness of cross-over locations, we expect substantial variation around the expected mean HI, even in the absence of BDMIs.

To estimate the amount of introgression in the BC_2_ pools, we computed the HI in both pools along the genome based on species-diagnostic SNPs (variants that are differentially fixed between the parental pools). Differentially fixed SNPs were defined as SNPs with allele frequency 1 in one parental pool and 0 in the other one (1 = all reads supporting the alternate allele, 0 = all reads supporting the reference allele). The total number of SNPs that were differentially fixed between the parental species was 1,668,294 when using *D. flavomontana* reference genome, and 1,570,556 when using *D. montana* reference genome. For each differentially fixed SNP between the species, allele frequencies were calculated by dividing “alternate read depth (AD)” by “the total read depth (DP)”. To enable comparison between backcross directions, the allele frequencies for non-reference alleles were calculated with the formula “1 - allele frequency” (e.g. allele frequency of 87.5% would become 12.5%). Finally, given that a maximum allele frequency for a SNP in a hybrid is 0.5, any SNPs with an allele frequency over 0.5 were discarded (78 out of 1,668,372 and 48 out of 1,570,604 when using *D. flavomontana* and *D. montana* reference genomes, respectively).

We compared colinear and inverted parts within each chromosome in terms of the density of diagnostic SNPs. Each chromosome was divided into 200kb non-overlapping windows and the number of diagnostic SNPs in each window was counted using a custom script (https://github.com/vihoikka/SNP_mapper/blob/main/snp_binner.py). When analysing data using *D. flavomontana* reference genome, the chromosomes were divided in 53-153 windows depending on the chromosome length, while the respective values for *D. montana* reference genome were 55-163 windows per chromosome. The data was analysed using a generalised linear model (*glm* function) with a Poisson distribution, where the number of window-wise SNPs was used as a response variable, and either different chromosomes, or different genomic partitions (colinear, inverted) within each chromosome were used as explanatory variables. The analyses were performed in base R using R v1.2.1335-1 and R studio v3.6.1.

Using the diagnostic SNPs, we calculated the mean hybrid index (HI) and its standard deviation separately for different chromosomes for BC_2_fla and BC_2_mon pools. We also calculated the number of SNPs without any introgressed material (HI = 0%) separately for each chromosome for both pools. Finally, we plotted HI in non-overlapping windows of 400 SNPs for each chromosome and BC_2_ pool using a custom script (https://github.com/vihoikka/SNP_mapper/blob/main/datasmoother2.py). In principle, crossover (CO) events involving the two ancestral backgrounds (Fisher junctions; Fisher, 1954) should be visible as step changes in the HI of each pool. Assuming on average one CO per chromosome and female meiosis, the expected number of CO events per chromosome generated during the experiment is given by the total number of females (nBC_1_ + nBC_2_; Table S3) contributing to each pool (96 and 104 for BC_2_mon and BC_2_fla pools, respectively). Note that the number of Fisher junctions between *D. montana* and *D. flavomontana* ancestral material is lower since not all CO events in BC_1_ females generate junctions between heterospecific ancestry. In practice, however, the resolution especially for the junctions that are unique to a single BC_2_ individual (which correspond to a change in allele frequency of 1/82) is limited by the randomness in sequencing coverage of the pool.

Given that this experiment was started with a single-pair cross between the parental species and continued with repeated backcrosses between hybrid females and parental males, all backcross individuals inherited a maximum of one allele per locus from the donor species (Fig. 1). Thus, the genomes of BC individuals are a mosaic of two types of tracts: i) homozygous for the genetic background of the recipient species or ii) heterozygous between species. This limits the types of BDMIs that can be expressed (Table 1). Dominant-dominant pairwise BDMIs arise already in the F_1_ generation and, if severe, can cause sterility/inviability in both sexes. Recessive-recessive pairwise BDMIs cannot be detected in our experiment even if they were X-linked since i) all BC individuals involved in the experiment were females (no hemizygosity), and ii) the expression of these incompatibilities would require homozygous tracts for both species (Fig. 1). Hence, dominant-recessive BDMIs are the only strong postzygotic barriers that we expect to detect in this study.

### Simulating the backcross and re-sequence experiment

Given the stochastic nature of inheritance of chromatids in gametes and the randomness of cross-over locations in meiosis, we expected substantial variation in the mean HI (in the BC_2_ pools for each chromosome) around the expectation of 12.5% (Fig. 1). To evaluate whether the observed mean HI of each chromosome deviates significantly from that expected under simple models of introgression with or without inversions and/or extreme BDMIs, we simulated the crossing experiment under three different scenarios using *Mathematica* (Wolfram Research, Inc., version 11.02 Champaign, IL). All simulations were conditioned on the number of BC_2_ females each BC_1_ female contributes to the pool (Table S3). We also assumed one cross-over per female per chromosome in meiosis (a map length of 50cM). Given that the experiment involves two generations of crosses between hybrid females and pure parental males, our simulation only tracks the haplotype of female gametes contributing to BC_1_ and BC_2_ individuals. All *in silico* backcross experiments were simulated, separately for each chromosome, 10,000 times to obtain 5% and 95% quantiles for the mean HI.

First, we simulated the experiment under a simple null model of neutral introgression, i.e. assuming no BDMIs and no cross-over suppression due to inversions (SIM1, Fig. 2A). Second, we simulated the experiment similarly under neutrality, but including the breakpoint locations of inversions that are alternately fixed between *D. montana* and *D. flavomontana*. This was done simply by disallowing cross-over events within inverted regions (inversions breakpoints in Table S2), i.e. we did not attempt to include interchromosomal effects (SIM2, Fig. 2B). Third, we simulated the experiment under a model that assumes a single BDMI at a random position within the inverted part of the chromosome (SIM3, Fig. 2C). This single locus cannot be introgressed beyond the F_1_ generation, i.e. BC_1_ and BC_2_ females that are heterozygous for this locus are not produced. Note that while we refer to this as a BDMI for simplicity, we did not explicitly simulate pairwise incompatibilities. Thus, this locus can be regarded as a BDMI involving a dominant allele on the introgressing background (donor species) that is incompatible with one or more recessive alleles in the recipient background.

**Figure 2.**
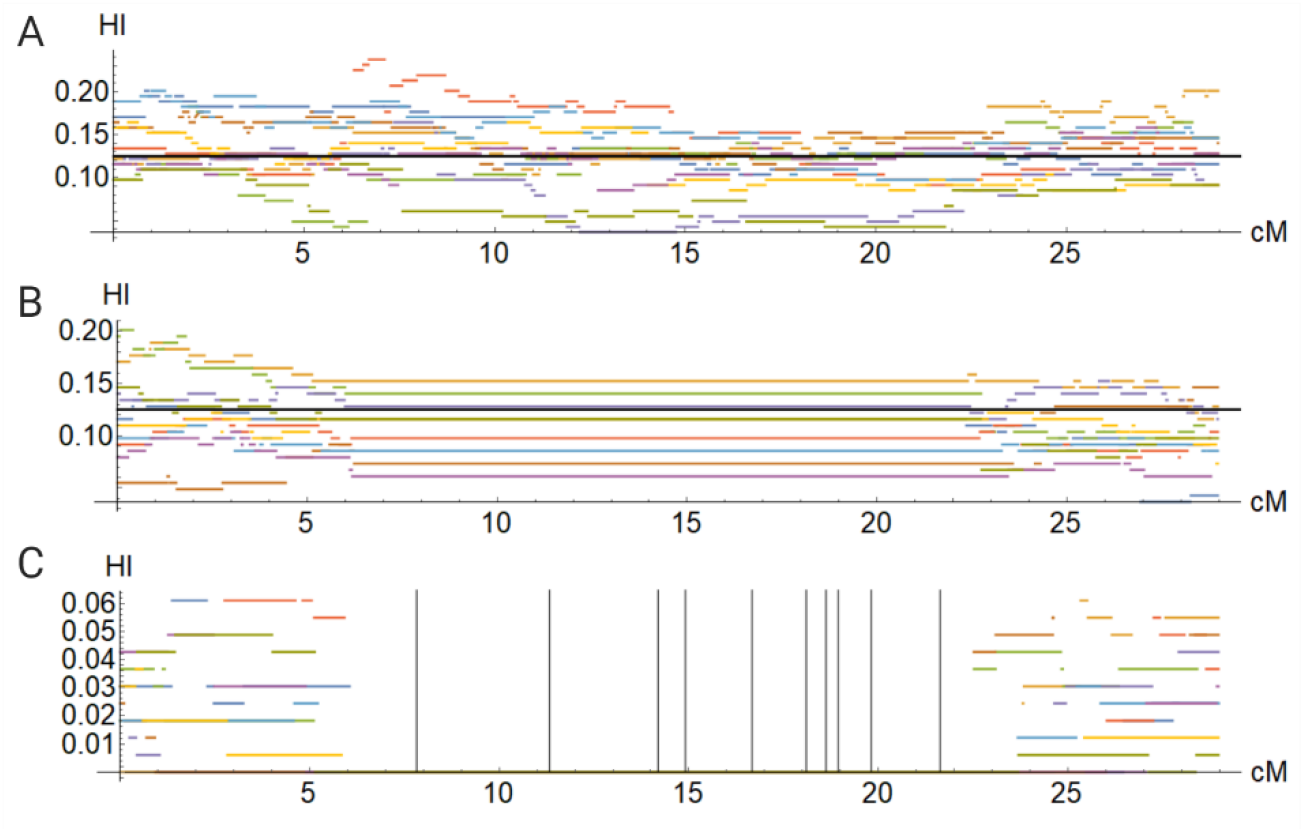
Introgression experiment was simulated under different scenarios. Example plots of simulated hybrid indices (HI) (A) under neutrality, (B) in the presence of neutral inversions, and (C) in the presence of inversions with a single dominant BDMI (grey vertical lines illustrate BDMIs). For simplicity, here simulations were run 10 times.

## Results

### BC_1_ females from the backcrosses towards *D. flavomontana* showed stronger genetic incompatibilities / postzygotic isolation than the ones from the backcrosses towards *D. montana*

In BC_1_ generation, the proportion of fertile females was 75% and 42% among the BC_1_mon and BC_1_fla hybrids, respectively, and was significantly reduced in both reciprocal crosses when compared to the expected fertility of 1 (BC_1_mon: t_19_ = −2.52, P = 0.021; BC_1_fla: t_54_ = −8.67, P = 8.371e^-12^). Furthermore, the proportion of fertile BC_1_mon females (75%) was significantly higher than that of BC_1_fla females (42%) (GLM, z_1,73_= −2.45, P = 0.015; Fig. S2). These findings show that while both crosses suffer from BDMIs affecting female fertility, these incompatibilities are more pronounced in backcrosses towards *D. flavomontana* than towards *D. montana* (asymmetric postzygotic isolation, or unidirectional incompatibilities in the sense of Turelli & Moyle, 2007).

### Genetic divergence between *D. montana* and *D. flavomontana* has accumulated within inverted chromosome regions especially on the X chromosome

We performed all genomic analyses using both *D. flavomontana* and *D. montana* reference genomes to be able to evaluate the potential effect of reference bias on the results. Here, we focus mainly on analyses that use *D. flavomontana* as a reference genome, since the backcrosses towards *D. flavomontana* showed more evidence for incompatibilities than the ones towards *D. montana*. Results based on the *D. montana* reference genome are also discussed here, but the corresponding figures and tables are given in the Supporting information.

Irrespective of which species was used as a reference genome, the density of SNPs that were differentially fixed between *D. montana* and *D. flavomontana* parental pools was higher on the X chromosome than on any of the autosomes (P < 0.001; Fig. 3; Fig. S3; Table S4). Moreover, the density of fixed differences was higher in inverted compared to the colinear regions within each chromosome containing inversions (P < 0.001; Fig. 3; Fig. S3; Table S5), as expected due to the reduction in recombination within inverted regions (note that chromosomes 2R and 3 have no inversions).

**Figure 3.**
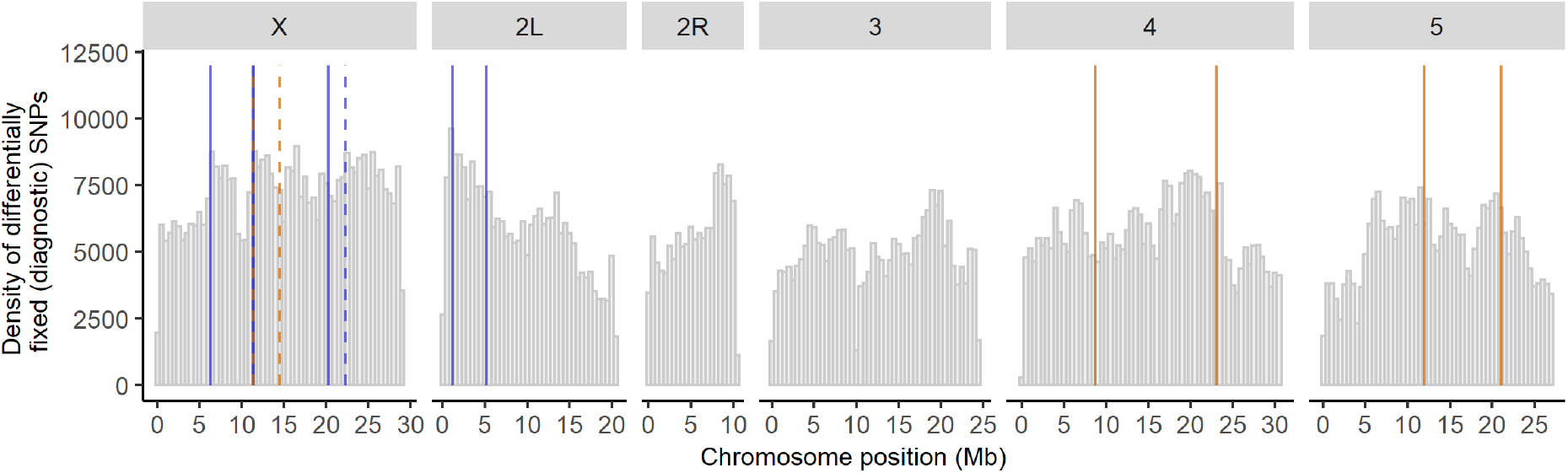
Density of differentially fixed SNPs (in 200kb windows) between the parental species across each chromosome (*D. flavomontana* used as a reference genome). Orange and blue vertical lines represent species-specific *D. flavomontana* and *D. montana* chromosomal inversions, respectively. Solid and dashed vertical lines describe breakpoints of different inversions. Chromosome 2 involves left (2L) and right (2R) arms separated by a submetacentric centromere. Corresponding data using *D. montana* as the reference genome shown in Fig. S3.

### Large differences in HI between chromosomes – evidence for BDMIs located within X chromosomal inversions

The mean amount of introgression (hybrid index, HI) of hybrids backcrossed to *D. montana* (BC_2_mon) did not deviate significantly from the neutral expectation of 12.5% for any chromosome (SIM1). This was true irrespective of whether the reference genome of *D. flavomontana* (Fig. 4, 5A, S4; Table 6) or *D. montana* (Fig. S5, S6, S7A, Table 6) was used. Moreover, in both analyses, the fraction of diagnostic SNPs that showed no introgression (HI = 0 in the BC_2_mon pool) was low (0.02-0.20% and 0.03-0.29% depending on whether the *D. flavomontana* or *D. montana* genome was used as a reference), across the entire genome (Table S6).

**Figure 4.**
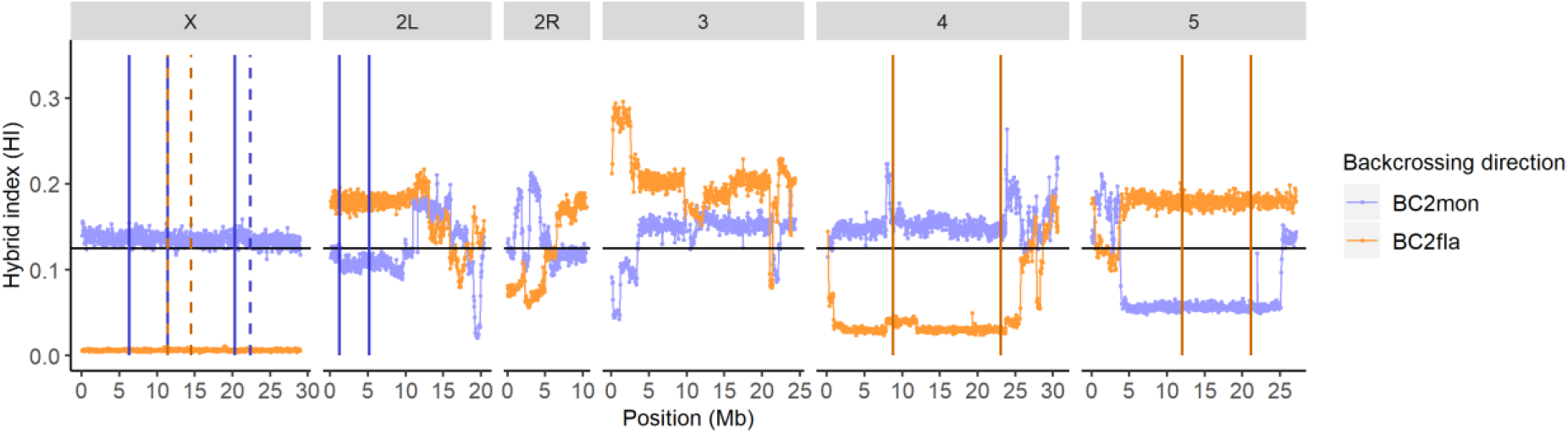
Observed hybrid index (HI) of 2^nd^ backcross generation female pools towards *D. montana* (BC_2_smon) and *D. flavomontana* (BC_2_fla) in windows of 400 non-overlapping SNPs along the genome. The data is illustrated using the *D. flavomontana* reference genome. For chromosome 2 the left (2L) and right (2R) arms are separated by a metacentric centromere. The black horizontal line represents the expected amount of introgression, HI = 12.5 %, under neutrality. Vertical lines represent species-specific *D. flavomontana* (yellow) and *D. montana* (blue) chromosomal inversions. Solid and dashed vertical lines show breakpoints of different inversions. Corresponding data using *D. montana* as the reference genome shown in Fig. S5.

**Figure 5.**
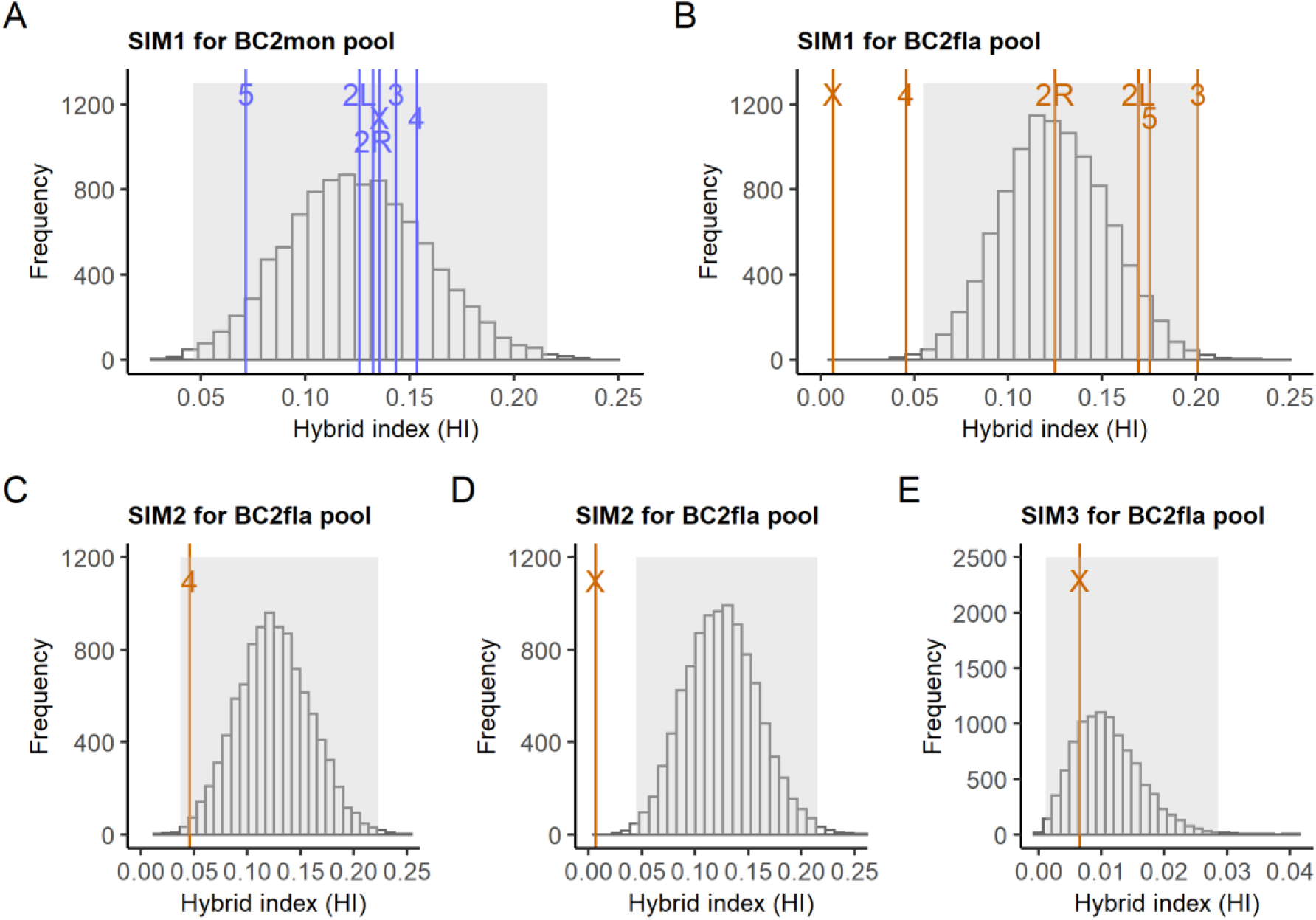
Hierarchical representation of the most meaningful simulations (10,000 replicates/simulation) of the 2^nd^ generation backcross experiments towards *D. montana* (BC_2_mon) and *D. flavomontana* (BC_2_fla) (*D. flavomontana* was used as a reference genome). The grey area of each figure represents Bonferroni corrected 5% and 95% quantiles and the space between them. We consider a mean HI outside of this range statistically different from the simulated model. Simulations under neutrality (SIM1) and the observed mean hybrid index (HI) of each chromosome for (A) BC_2_mon pool and (B) BC_2_fla pool. Simulations under neutral inversions (SIM2) and observed mean HI of BC_2_fla pool for (C) the 4^th^ chromosome and (D) the X chromosome. (E) Simulations involving inversions with a single locus against introgression (SIM3) and observed mean HI for the X chromosome of BC_2_fla pool. Corresponding data using *D. montana* as the reference genome shown in Fig. S7.

In contrast, BC_2_fla hybrids showed a significant reduction in mean HI compared to the neutral scenario (SIM1) for the 4^th^ and the X chromosome, and these results were again robust to the choice of reference genome (*D. flavomontana* genome: Fig. 4, 5B, S4, Table S6; *D. montana* genome: Fig. S5, S6, S7B, Table S6). Interestingly, and irrespective of which reference genome was used, the reduced introgression on the 4^th^ chromosome could be explained by the reduction in cross-over rate due to inversion present on this chromosome, without invoking any selection acting on incompatibilities (SIM2) (*D. flavomontana* genome: Fig. 4, 5C, S4; *D. montana* genome: Fig. S5, S6, S7D). Under this scenario, the mean HI showed no deviation from the expectation of 12.5% under neutrality but had an increased variance across simulation replicates.

In contrast to the pattern of the chromosome 4, the observed decrease in mean HI of BC_2_fla hybrids on the X chromosome could not be explained solely by a reduction in cross-over rate due to inversions (SIM2) (*D. flavomontana* genome: Fig. 4, 5D, S4; *D. montana* genome: Fig. S5, S6, S7F). Instead, our simulations show that the drastic reduction in mean HI on the X chromosome is compatible with a single or multiple dominant incompatibility locus/loci residing within the X inversions (SIM3) (*D. flavomontana* genome: Fig. 4, 5E, S4; *D. montana* genome: Fig. S5, S6, S7G). In other words, the data are consistent with at least one dominant X chromosomal *D. montana* allele that interacts negatively with autosomal homozygous recessive *D. flavomontana* alleles. Intriguingly, depending on the reference genome used, 39.4-44.5% of the differentially fixed SNPs between the species on the X chromosome showed no introgression, emphasising the strength of the X-effect (Table S6). For the autosomes, the fraction of diagnostic SNPs that showed no introgression into *D. flavomontana* varied from 0.14% to 2.58%, depending on the chromosome and the choice of reference genome.

Chromosome 3 and 5 showed an increased HI in the BC_2_fla pool relative to the neutral expectation of 12.5% (SIM1; Fig. S5-S6; Fig. S7B). However, the interpretation of this finding depends on the choice of reference genome. Using *D. flavomontana* as a reference genome (which likely underestimates introgression of *D. montana* alleles into the BC_2_fla pool), the estimated mean HI for the chromosomes 3 and 5 were within the 95^th^ percentile for the neutral case (SIM1; Fig. 4; 5B, S4). However, when we used *D. montana* as a reference genome (which likely overestimates introgression of *D. montana* alleles into the BC_2_fla pool), BC_2_fla hybrids showed a significant increase in mean HI relative to the neutral scenario (SIM1) for both chromosomes (Fig. S5, S6, S7B). In this case, we find that the increase in introgression on the 5^th^ chromosome was compatible with a reduction in cross-over rate due to the inversion present on this chromosome, without invoking any selection acting on incompatibilities (SIM2) (Fig. S7E). In contrast, the mean estimated HI in BC_2_fla hybrids for chromosome 3 (which has no known inversion differences between the two species) was not compatible with any of the simple scenarios we simulated. Given that we have either assumed neutrality or a single dominant incompatibility locus, which is maximally deleterious, this is perhaps unsurprising (see Discussion).

## Discussion

A major theme in speciation research is to understand how the loci inducing genetic incompatibilities (BDMIs) in interspecific crosses are distributed across the genome, what role chromosomal inversions and the X chromosome may play in their distribution and what types of epistatic interactions matter for BDMIs (reviewed in (Coughlan & Matute, 2020; Coyne, 2018; Faria, Johannesson, Butlin, & Westram, 2018; Hoffmann & Rieseberg, 2008)). To shed light on these questions, we performed reciprocal backcrosses between *D. montana* and *D. flavomontana* and traced the regions of reduced introgression in 2^nd^ backcross generation (BC_2_) females.

### Postzygotic barriers between *D. montana* and *D. flavomontana* show asymmetry in their strength

We have previously shown that pre- and postzygotic barriers between *D. montana* females and *D. flavomontana* males are practically complete, while both types of barriers between *D. flavomontana* females and *D. montana* males are weaker (Poikela et al., 2019). In crosses between *D. flavomontana* females and *D. montana* males, F_1_ hybrid males are sterile, but roughly half of the F_1_ females are fertile (Noora Poikela et al., 2019). Accordingly, here we backcrossed fertile F_1_ females with the males of both parental species, and observed a clear asymmetry in the strength of postzygotic barriers between the two backcross directions. BC_1_ hybrid females born from the backcrosses between F_1_ females and *D. montana* males showed rather high fertility, and the genetic incompatibilities in BC_2_ females had no detectable effect. In contrast, when backcrossing F_1_ hybrid females with *D. flavomontana* males, more than half of the BC_1_ females were sterile, and BC_2_ females showed signs of strong BDMIs. This asymmetry could be a consequence of a history of unidirectional introgression from *D. montana* into *D. flavomontana* in nature (Poikela et al., 2022), if it had induced selection against introgression at certain loci especially within the X chromosomal inversions, but homogenised genetic divergence on colinear regions. This kind of pattern in the permeability of species boundaries has been found to contribute to speciation also in other species (Harrison & Larson, 2014).

It is surprising that introgression has not occurred from *D. flavomontana* to *D. montana* in nature, given that backcrossing towards *D. montana* (BC_2_mon) was relatively successful in this study. The most obvious reason for this discrepancy is that laboratory experiments may not reveal all reproductive barriers relevant in wild populations. For example, hybrids may have problems in mate choice in the wild, or they may face challenges to feed or reproduce on species-specific host trees. Moreover, also the male hybrids regain fertility in backcross generations (data not shown), which may contribute to introgression in nature. Finally, BDMIs may well be stronger between *D. montana and D. flavomontana* populations living in close contact.

### The role of inversions and the X chromosome in reducing recombination and introgression from *D. montana* to *D. flavomontana* (BC_2_fla pool)

Inversions have been suggested to contribute to speciation, when three criteria are met: closely related species must carry alternatively fixed inversions, the inversions suppress recombination, and this suppression of recombination facilitates reproductive isolation (Faria & Navarro, 2010). *D. montana* populations on different continents are known to have a high number of fixed and polymorphic inversions (Morales-hojas, Päällysaho, Vieira, Hoikkala, & Vieira, 2007; Throckmorton, 1982), while there is less data on *D. flavomontana* inversions (Throckmorton, 1982). Using long- and short-read genomic data, we have recently identified several alternatively fixed inversions in *D. montana* and *D. flavomontana* across species’ distribution in North America, and shown that these inversions have increased genetic divergence and lower historical introgression compared to colinear chromosome regions (Poikela et al., 2022). In the present study, we show that these inversions have an increased number of alternatively fixed SNPs compared to colinear regions, which is in agreement with their increased genetic divergence shown in Poikela et al. (2022). We have also shown that large swathes of species-specific ancestry are retained within inverted chromosome regions (Fig. 4), which suggests that inversions effectively suppress recombination in early backcross hybrids. Finally, we find that the drastic reduction in introgression on the X chromosome can be explained by inversions that are associated with at least one dominant X chromosomal *D. montana* incompatibility allele interacting negatively with recessive autosomal *D. flavomontana* alleles. This negative epistatic interaction could cause the observed low hybrid fertility, and supports the idea that inversions act as strong barriers to gene flow by facilitating the establishment of BDMIs (Hoffmann & Rieseberg, 2008; Navarro & Barton, 2003; Noor et al., 2001).

While the involvement of the X chromosome in hybrid problems may not be surprising (see e.g. Masly & Presgraves, 2007; Tao, Chen, Hartl, & Laurie, 2003), the fact that it involves a dominant incompatibility locus is. The “dominance theory” (e.g. Turelli & Orr, 1995, 2000), which aims to explain the disproportionate role of the X chromosome in hybrid incompatibilities, relies on the presence of recessive incompatibilities on the X and therefore cannot explain our result. However, the “dominance theory”, as well as the “faster-male theory” and dosage compensation (reviewed in Coyne, 2018; Presgraves, 2008), can still explain the hybrid male sterility previously observed in crosses between *D. flavomontana* and *D. montana* (Poikela et al., 2019). Accumulation of meiotic drive elements on the X chromosome could be another plausible explanation for the large X-effect in general (reviewed in Patten, 2018), but this is unlikely in our system as the meiotic drive systems described in *Drosophila* are typically involved in sperm killing and not in female sterility (Courret et al., 2019). Although cytoplasmic incompatibilities have been detected in other *montana* complex species of the *Drosophila virilis* group (Patterson, 1952; Throckmorton, 1982), they are not likely to play a major role in these crosses since all hybrids had *D. flavomontana* cytoplasm (and crosses were more unsuccessful in this direction). Finally, the large X-effect we detected in the present study could potentially be explained by “faster X evolution”, based on the idea that selection increases the frequency of advantageous recessive alleles more effectively on the X chromosome than on autosomes, irrespectively of whether the incompatibilities themselves are recessive (Charlesworth et al. 1987, 2018). Also, the X chromosome could simply contain more genes that are prone to create postzygotic isolation than those on the autosomes (Coyne, 2018).

Several autosomes showed deviations from the expected hybrid indices in the BC_2_fla pool. Based on our simulations, the reduced introgression on the 4^th^ chromosome could be explained by inversions’ ability to restrict recombination which increases the variance in chromosome-wide HI. However, if we calculate the expected allele frequencies for a dominant–recessive BDMI by hand for the first two backcross generations, the allele frequencies (i.e. HI) after selection would be 1/22 (4.5%) for the dominant and 2/11 (18.2%) for the recessive *D. montana* allele in the BC_2_fla pool (see Fig. S8). These frequencies are close to the observed frequencies e.g. on chromosomes 4 (4.6%) and 5 (17.5%), respectively. It is therefore tempting to speculate that pairwise BDMI loci could exist on these chromosomes. Finally, chromosomes 3 and 5 showed increased introgression in the BC_2_fla pool, but only in analyses using *D. montana* as a reference. This effect may be due to an overestimation of *D. montana* alleles in the BC_2_fla pool (i.e. reference bias). Alternatively, the increased introgression on 5^th^ chromosome could be explained by inversions’ ability to restrict recombination, increasing the variance in chromosome-wide HI. However, the drastic increase in introgression on the 3^rd^ chromosome, which lacks species-specific inversions, was not explained by any of our simulations. We note that our simulations did not consider an interchromosomal effect, where inversions may trigger an increase in recombination on other freely recombining chromosomes (Crown et al., 2018; Stevison, Hoehn, & Noor, 2011). However, this would only decrease the variance in HI on chromosomes lacking fixed inversions and, and thus it cannot explain the increase in HI for chromosome 3 in the BC_2_fla pool.

In future research, combining the crosses with quantitative trait loci (QTL) analyses might help to link BDMIs to e.g. specific genes (Johnson, 2010), gene duplicates or transposons (Bikard et al., 2009; Masly, Jones, Noor, Locke, & Orr, 2006). BDMI genes could also be searched by tracing whole-genome gene expression data in interspecific hybrids (Satokangas, Martin, Helanterä, Saramäki, & Kulmuni, 2020). However, recombination suppression of inversions presents a challenge for mapping BDMIs, and would in theory require a complex reversion of the X chromosomal inversions with genome editing tools, and repeating the current experiment to narrow down the regions of reduced introgression (Hopkins, Tyukmaeva, Gompert, Feder, & Nosil, 2020). Overall, finding the exact loci driving species’ isolation may be difficult, as BDMIs are often complex and co-evolve with rapidly evolving heterochromatic DNA (Satyaki et al., 2014).

## Conclusions

“Introgress-and-resequence” studies that combine interspecific backcrosses with genome-wide analyses and simulations are an effective approach for identifying BDMIs, in particular those involving dominant alleles. Our study supports the idea that inversions aid the accumulation of BDMIs due to reduced recombination, and shows that strong BDMIs coupled with suppressed recombination effectively restrict introgression beyond the inverted part of the genome in the first two backcross generations. We conclude that the large X-effect we observed in our experiment may result from at least one dominant incompatibility locus residing within several overlapping inversions. If the design were extended to study interspecific F_2_ hybrids, assuming that the F_1_ female and male hybrids are viable and fertile, one could investigate recessive-recessive BDMIs in the same way. Overall, we provide a novel framework for investigating the role of inversions and the X chromosome as genetic barriers to introgression, which we hope will encourage similar studies on a larger number of species and strains.

## Acknowledgements

This work was supported by the grants from Academy of Finland project 267244 to AH and projects 268214 and 322980 to MK, as well as a grant from Emil Aaltonen to NP. KL was supported by a Natural Environmental Research Council (NERC) UK Independent Research fellowship (NE/L011522/1). We thank Anna-Lotta Hiillos for performing DNA extractions and Ville Hoikkala for his help with data analysis.

## Data Accessibility and Benefit-Sharing

Raw reads will be made publicly available in SRA and other data (phenotypic and allele frequency data, reference genomes for both species, *Mathematica* notebooks including simulations, and Unix and R commands) in Dryad at the time of publication.

## Author Contributions

KL, AH and NP designed the study. NP performed the hybrid backcrosses and analysed the genomic data with input from KL and DRL. KL performed the simulations. AH and MK supervised and funded the research. NP, AH and KL drafted the manuscript and all authors finalised it.

## Ethics declaration

Neither species is endangered, and the flies were collected along watersides on public lands outside National and State parks, where insect collecting does not require permits in the USA (The Wilderness Act of 1964, section 6302.15).

