## Supporting information for "Experimental introgression in *Drosophila*: asymmetric postzygotic isolation associated with chromosomal inversions and an incompatibility locus on the X chromosome"

Supplementary Tables ………………………………………………………………………. 1
Supplementary Figures ……………………………………………………………………. 5
Supplementary References ………………………………………………………………. 9

### Supplementary Tables:

Table S1. Number and mean length of Illumina paired-end reads of each pool before and after filtering, as well as read depth (coverage) after mapping to *D. montana* and *D. flavomontana* reference genomes.

| | | Reads (in millions) | | Mean length (bp) of forward and reverse reads | | Mean coverage after mapping to | |
|---|---|---|---|---|---|---|---|
| **Pool** | **Lane** | **Before filtering** | **After filtering** | **Before filtering** | **After filtering** | ***D. flavomontana* genome** | ***D. montana* genome** |
| *D. montana* pool | lane1 | 160.2 | 156.3 | 150, 150 | 142, 141 | 192.7 | 204.1 |
| | lane2 | 160.7 | 156.7 | 150, 150 | 143, 142 | | |
| *D. flavomontana* pool | lane1 | 169.8 | 164.6 | 150, 150 | 143, 141 | 163.6 | 152.8 |
| | lane2 | 170.3 | 165.0 | 150, 150 | 143, 142 | | |
| BC2*mon* pool | lane1 | 177.8 | 173.3 | 150, 150 | 143, 142 | 169.3 | 178.8 |
| | lane2 | 178.0 | 173.5 | 150, 150 | 143, 142 | | |
| BC2*fla* pool | lane1 | 158.0 | 152.7 | 150, 150 | 143, 142 | 162.8 | 150.7 |
| | lane2 | 158.1 | 152.7 | 150, 150 | 143, 142 | | |

Table S2. Genomic coordinates of chromosomal inversions fixed between *D. montana* and *D. flavomontana* (Poikela et al., in prep.). Presence of inversions was checked from parental female pools, and they are illustrated against *D. flavomontana* and *D. montana* reference genomes.

| | | Inversion breakpoints in *D. flavomontana* chromosome-level genome | | | | | Inversion breakpoints in *D. montana* chromosome-level genome | | | | |
|---|---|---|---|---|---|---|---|---|---|---|---|
| **Chromosome** | **Species** | **Scaffold** | **Scaffold length** | **Start** | **End** | **Size (bp)** | **Scaffold** | **Scaffold length** | **Start** | **End** | **Size (bp)** |
| X | *D. montana* | X_chromosome | 28975156 | 6309450 | 20284519 | 13975070 | X_chromosome | 29140820 | 3994118 | 11174373 | 7180256 |
| X | *D. montana* | X_chromosome | 28975156 | 11367083 | 22321964 | 10954882 | X_chromosome | 29140820 | 6041058 | 20203141 | 14162084 |
| X | *D. flavomontana* | X_chromosome | 28975156 | 11368055 | 14487633 | 3119579 | X_chromosome | 29140820 | 17044656 | 20200540 | 3155884 |
| 2L | *D. montana* | 2L_chromosome | 20392248 | 1181134 | 5145967 | 3964834 | 2L_chromosome | 20245637 | 15098446 | 19046306 | 3947861 |
| 4 | *D. flavomontana* | 4_chromosome | 30698533 | 7855585 | 23745872 | 15890288 | 4_chromosome | 32544174 | 3246708 | 17382172 | 14135465 |
| 5 | *D. flavomontana* | 5_chromosome | 27217941 | 11949884 | 21134310 | 9184427 | 5_chromosome | 26508887 | 9422591 | 18649749 | 9227159 |

Table S3. Fertile and viable $F_1$ and $BC_1$ females, produced by a single-pair cross of *D. flavomontana* female and *D. montana* male, contributed to the production of the sequenced $BC_2$ females. These numbers were used to repeat the experiment *in silico* (simulations).

| **Backcrossed to *D. montana* (BC2mon)** | | |
|---|---|---|
| **F1-generation** | **BC1-generation** | **BC2 generation** |
| F1♀xmon♂ | BC1♀ x mon♂ | 16 |
| F1♀xmon♂ | BC1♀ x mon♂ | 3 |
| | BC1♀ x mon♂ | 6 |
| | BC1♀ x mon♂ | 10 |
| F1♀xmon♂ | BC1♀ x mon♂ | 1 |
| | BC1♀ x mon♂ | 3 |
| | BC1♀ x mon♂ | 2 |
| | BC1♀ x mon♂ | 2 |
| | BC1♀ x mon♂ | 5 |
| F1♀xmon♂ | BC1♀ x mon♂ | 8 |
| | BC1♀ x mon♂ | 6 |
| | BC1♀ x mon♂ | 4 |
| | BC1♀ x mon♂ | 9 |
| | BC1♀ x mon♂ | 7 |
| | | tot. 82 females |
| **Backcrossed to *D. flavomontana* (BC2fla)** | | |
| **F1 generation** | **BC1 generation** | **BC2 generation** |
| F1♀xfla♂ | BC1♀ x fla♂ | 4 |
| | BC1♀ x fla♂ | 1 |
| | BC1♀ x fla♂ | 5 |
| F1♀xfla♂ | BC1♀ x fla♂ | 9 |
| | BC1♀ x fla♂ | 5 |
| | BC1♀ x fla♂ | 1 |
| | BC1♀ x fla♂ | 3 |
| | BC1♀ x fla♂ | 4 |
| | BC1♀ x fla♂ | 3 |
| | BC1♀ x fla♂ | 1 |
| F1♀xfla♂ | BC1♀ x fla♂ | 2 |
| | BC1♀ x fla♂ | 2 |
| | BC1♀ x fla♂ | 8 |
| | BC1♀ x fla♂ | 3 |
| | BC1♀ x fla♂ | 9 |
| | BC1♀ x fla♂ | 4 |
| | BC1♀ x fla♂ | 1 |
| | BC1♀ x fla♂ | 6 |
| | BC1♀ x fla♂ | 6 |
| | BC1♀ x fla♂ | 1 |
| | BC1♀ x fla♂ | 2 |
| | BC1♀ x fla♂ | 2 |
| | | tot. 82 females |

Table S4. Summary of the effect of chromosome on the number of differentially fixed SNPs between *D. montana* and *D. flavomontana*. Significant P-values are in bold. The data were analysed twice, using both *D. flavomontana* and *D. montana* reference genomes.

| **Analyses based on *D. flavomontana* reference genome** | **Model** | **z-statistic** | **Df** | **P-value** |
|---|---|---|---|---|
| | Intercept (X chromosome) | 5172.62 | 5,703 | < 0.001 |
| | chromosome 2L | -74.54 | | **< 0.001** |
| | chromosome 2R | -63.05 | | **< 0.001** |
| | chromosome 3 | -147.72 | | **< 0.001** |
| | chromosome 4 | -107.30 | | **< 0.001** |
| | chromosome 5 | -127.63 | | **< 0.001** |
| **Analyses based on *D. montana* reference genome** | **Model** | **z-statistic** | **Df** | **P-value** |
| | Intercept (X chromosome) | 4978.29 | 5,722 | < 0.001 |
| | chromosome 2L | -68.66 | | **< 0.001** |
| | chromosome 2R | -73.21 | | **< 0.001** |
| | chromosome 3 | -156.62 | | **< 0.001** |
| | chromosome 4 | -125.37 | | **< 0.001** |
| | chromosome 5 | -124.81 | | **< 0.001** |

Table S5. Summary of the effect of genomic region (collinear, inverted) on the number of differentially fixed SNPs between *D. montana* and *D. flavomontana*. Significant P-values are in bold. The data were analysed twice, using both *D. flavomontana* and *D. montana* reference genomes.

| **Analyses based on *D. flavomontana* reference genome** | **Chromosome** | **Model** | **z-statistic** | **Df** | **P-value** |
|---|---|---|---|---|---|
| | X | Intercept (colinear) | 3375.80 | 1,143 | < 0.001 |
| | | inverted | 23.40 | | **< 0.001** |
| | 2L | Intercept (colinear) | 3285.32 | 1,100 | < 0.001 |
| | | inverted | 76.08 | | **< 0.001** |
| | 4 | Intercept (colinear) | 3121.06 | 1,151 | < 0.001 |
| | | inverted | 62.73 | | **< 0.001** |
| | 5 | Intercept (colinear) | 3242.84 | 1,134 | < 0.001 |
| | | inverted | 40.72 | | **< 0.001** |
| **Analyses based on *D. montana* reference genome** | **Chromosome** | **Model** | **z-statistic** | **Df** | **P-value** |
| | X | Intercept (colinear) | 3163.79 | 1,144 | < 0.001 |
| | | inverted | 42.12 | | **< 0.001** |
| | 2L | Intercept (colinear) | 3139.21 | 1,99 | < 0.001 |
| | | inverted | 77.78 | | **< 0.001** |
| | 4 | Intercept (colinear) | 3235.59 | 1,161 | < 0.001 |
| | | inverted | 14.39 | | **< 0.001** |
| | 5 | Intercept (colinear) | 3052.98 | 1,131 | < 0.001 |
| | | inverted | 35.77 | | **< 0.001** |

Table S6. Information on the chromosome length (bp), the number and proportion (%) of differentially fixed SNPs (diagnostic SNPs) between parental species (*D. montana* and *D. flavomontana*) for different genomic regions, the observed mean hybrid index (HI) for different genomic regions and for both 2nd generation backcross pools (BC2mon and BC2fla), and the number and proportion (%) of SNPs that showed no introgression between species (HI=0). The data were analysed twice, using both *D. flavomontana* and *D. montana* reference genomes.

| **Analyses based on *D. flavomontana* as reference genome** | **Chromosome** | **Length (bp)** | **# and % of diagnostic SNPs between the species** | **Mean and SD of HI in BC2mon pool (%)** | **Mean and SD of HI in BC2fla pool (%)** | **# and % of SNPs with HI = 0 in BC2mon pool** | **# and % of SNPs with HI = 0 in BC2fla pool** |
|---|---|---|---|---|---|---|---|
| | **All** | 142,318,760 | 1,668,294 (1.17 %) | 12.76 ± 5.22 | 10.49 ± 8.85 | 1,880 (0.11 %) | 201,379 (12.07 %) |
| | **X** | 28,975,156 | 420,864 (1.45 %) | 13.55 ± 3.88 | 0.65 ± 0.76 | 74 (0.02 %) | 187,307 (44.51 %) |
| | **2L** | 20,392,248 | 244,886 (1.20 %) | 12.57 ± 4.81 | 16.92 ± 4.75 | 350 (0.14 %) | 474 (0.19 %) |
| | **2R** | 10,575,737 | 122,143 (1.15 %) | 13.24 ± 4.71 | 12.48 ± 5.71 | 23 (0.02 %) | 305 (0.25 %) |
| | **3** | 24,459,145 | 243,047 (0.99 %) | 14.33 ± 4.58 | 20.12 ± 5.79 | 315 (0.13 %) | 2,114 (0.87 %) |
| | **4** | 30,698,533 | 347,245 (1.13 %) | 15.33 ± 4.54 | 4.58 ± 4.17 | 698 (0.20 %) | 8,973 (2.58 %) |
| | **5** | 27,217,941 | 290,109 (1.07 %) | 7.14 ± 4.60 | 17.52 ± 4.72 | 420 (0.14 %) | 2,206 (0.76 %) |
| **Analyses based on *D. montana* as reference genome** | **Chromosome** | **Length (bp)** | **# and % of diagnostic SNPs between the species** | **Mean and SD of HI in BC2mon pool (%)** | **Mean and SD of HI in BC2fla pool (%)** | **# and % of SNPs with HI = 0 in BC2mon pool** | **# and % of SNPs with HI = 0 in BC2fla pool** |
| | **All** | 145,453,339 | 1,570,556 (1.08 %) | 10.99 ± 4.45 | 12.05 ± 9.92 | 2,754 (0.18 %) | 164,549 (10.45 %) |
| | **X** | 29,140,820 | 396,437 (1.36 %) | 11.55 ± 3.22 | 0.81 ± 0.88 | 104 (0.03 %) | 156,086 (39.37 %) |
| | **2L** | 20,245,637 | 229,022 (1.13 %) | 10.91 ± 4.15 | 19.23 ± 5.04 | 445 (0.19 %) | 330 (0.14 %) |
| | **2R** | 10,997,687 | 117,065 (1.06 %) | 11.68 ± 4.11 | 14.14 ± 6.27 | 25 (0.21 %) | 212 (0.18 %) |
| | **3** | 26,016,134 | 234,764 (0.90 %) | 12.41 ± 3.98 | 23.09 ± 5.73 | 620 (0.26 %) | 1,233 (0.53 %) |
| | **4** | 32,544,174 | 329,335 (1.01 %) | 13.09 ± 3.84 | 5.59 ± 4.92 | 951 (0.29 %) | 5,211 (1.58 %) |
| | **5** | 26,508,887 | 263,933 (1.00 %) | 6.03 ± 3.84 | 20.05 ± 4.93 | 609 (0.23 %) | 1,477 (0.56 %) |

### Supplementary Figures:

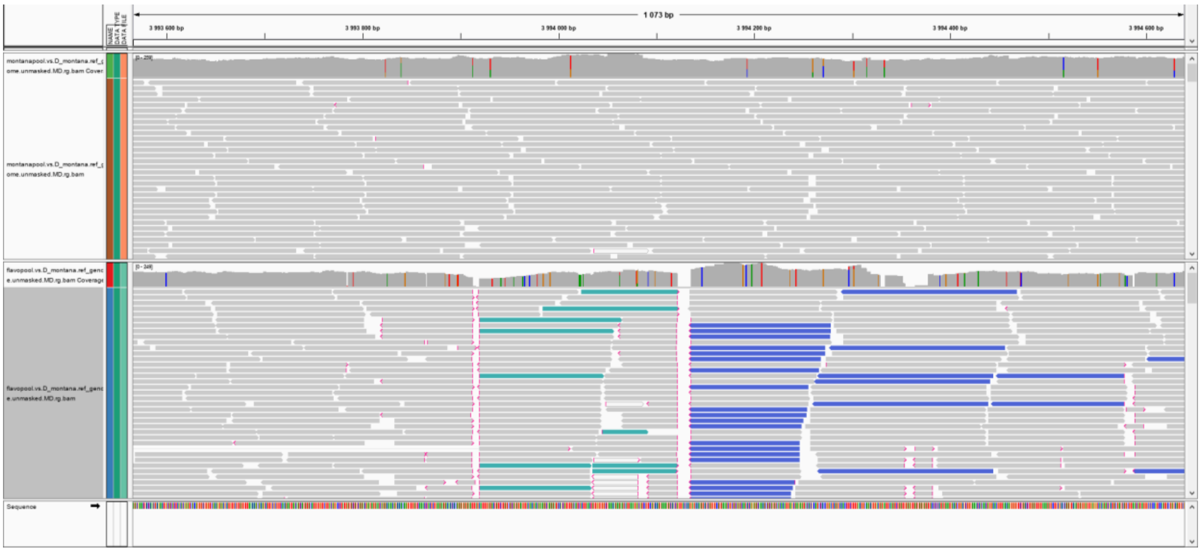

Figure S1. Example plot of an inversion breakpoint between *D. montana* and *D. flavomontana* illustrated with Interactive Genomics Viewer (Thorvaldsdóttir et al. 2012). Illumina reads of the parental strains (monSE13F37 above and flaMT13F11 below) are mapped against *D. montana* chromosome level reference genome. *D. montana* reads map well on the *D. montana* genome, while *D. flavomontana* reads split or are in reversed orientation. Red tip of a read represents the split of the read, and blue and green reads represent reads that are in reversed orientation relative to the reference genome.

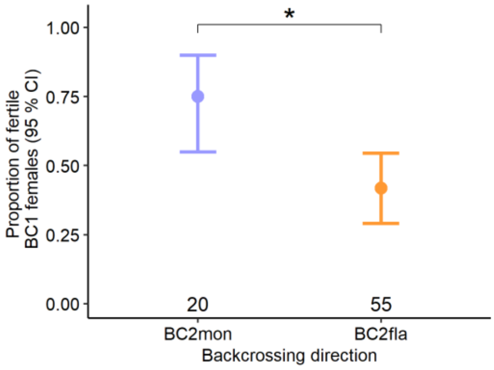


Figure S2. Proportion of fertile $BC_1$ females from backcrosses between $F_1$ females and *D. montana* males (blue) and between $F_1$ females and *D. flavomontana* males (orange). Numbers below error bars refer to the number of tested $BC_1$ females in each reciprocal cross. Error bars represent bootstrapped 95% confidence intervals (Mean ± CI).

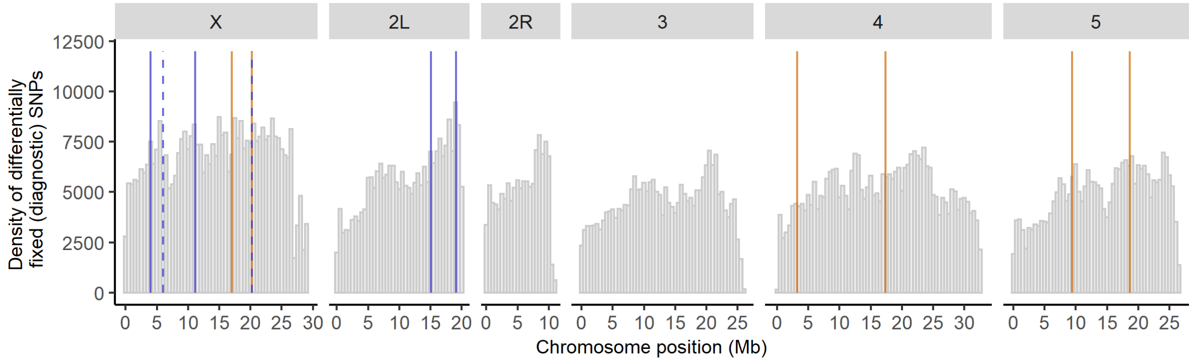


Figure S3. Density of differentially fixed SNPs (in 200kb windows) between parental species across each chromosome (*D. montana* used as a reference genome). Orange and blue vertical lines represent species-specific *D. flavomontana* and *D. montana* chromosomal inversions, respectively. Solid and dashed vertical lines describe breakpoints of different inversions. Chromosome 2 involves left (2L) and right (2R) arms separated by a submetacentric centromere.

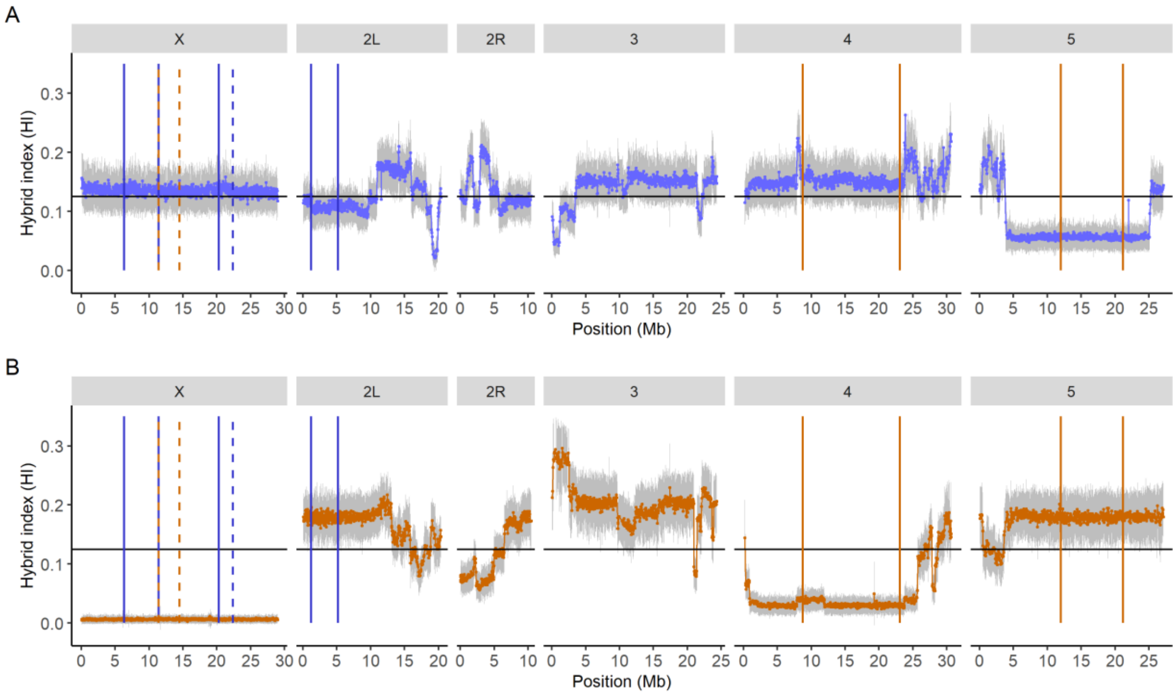


Figure S4. Observed hybrid index (HI) of 2nd backcross generation female pools towards (A) *D. montana* ($BC_2$mon) and (B) *D. flavomontana* ($BC_2$fla). The data was averaged over windows of 400 non-overlapping SNPs along the genome. The grey area shows the variation within each averaged data point measured as standard deviation. The data is illustrated using the *D. flavomontana* reference genome. The left (2L) and right (2R) arms of chromosome 2 are separated by a metacentric centromere. The black horizontal line represents the expected amount of introgression, HI = 12.5 %, under neutrality. Vertical lines represent species-specific *D. flavomontana* (orange) and *D. montana* (blue) chromosomal inversions. Solid and dashed vertical lines show breakpoints of different inversions.

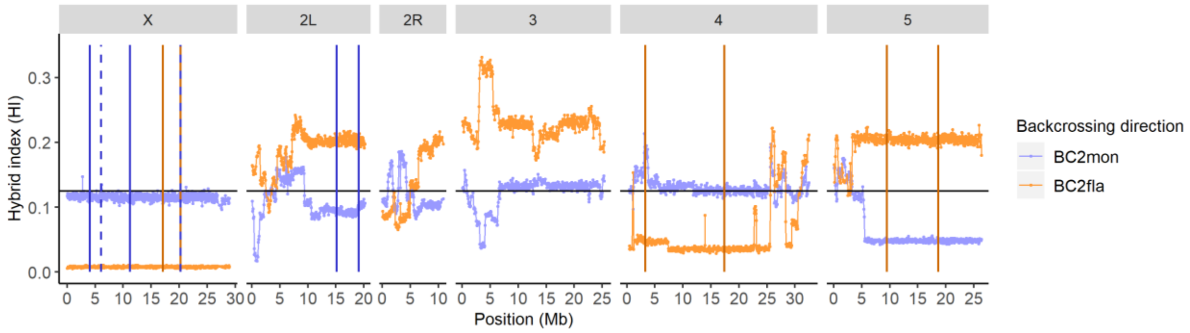


Figure S5. Observed hybrid index (HI) of 2nd backcross generation female pools towards *D. montana* ($BC_2$mon) and *D. flavomontana* ($BC_2$fla) in windows of 400 non-overlapping SNPs along the genome. The data is illustrated using the *D. montana* reference genome. For chromosome 2 the left (2L) and right (2R) arms are separated by a metacentric centromere. The black horizontal line represents the expected amount of introgression, HI = 12.5 %, under neutrality. Vertical lines represent species-specific *D. flavomontana* (orange) and *D. montana* (blue) chromosomal inversions. Solid and dashed vertical lines show breakpoints of different inversions.

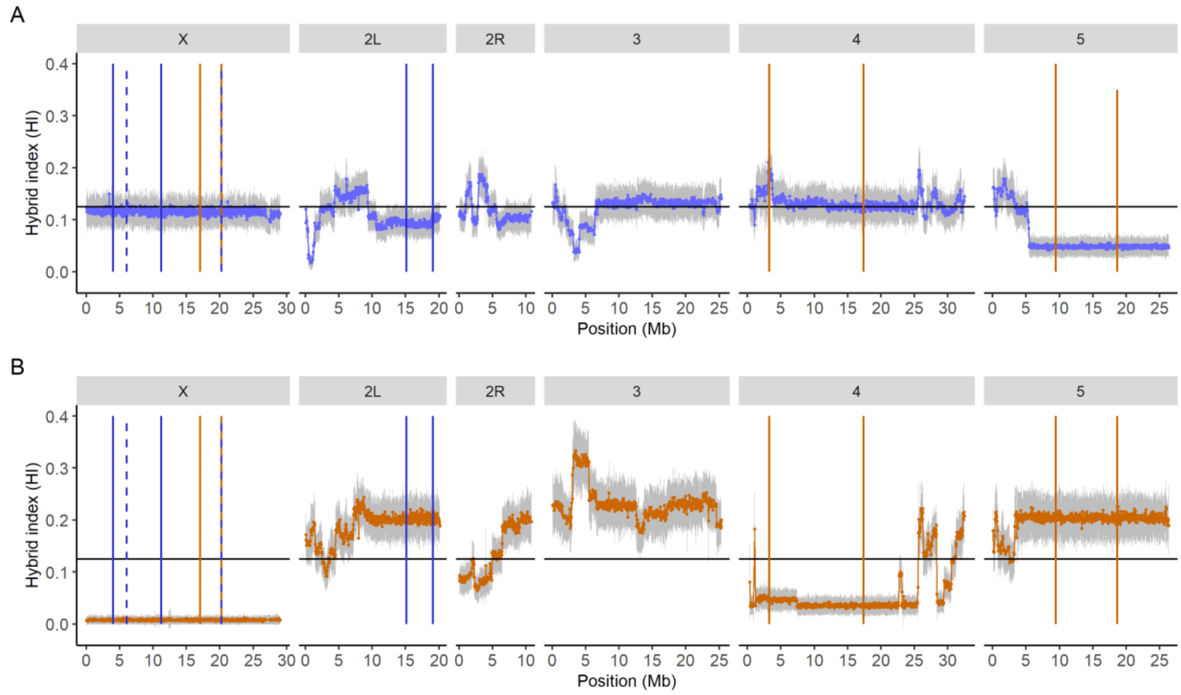


Figure S6. Observed hybrid index (HI) of 2nd backcross generation female pools towards (A) *D. montana* ($BC_2$mon) and (B) *D. flavomontana* ($BC_2$fla). The data was averaged over windows of 400 non-overlapping SNPs along the genome. The grey area shows the variation within each averaged data point measured as standard deviation. The data is illustrated using the *D. montana* reference genome. For chromosome 2 the left (2L) and right (2R) arms are separated by a metacentric centromere. The black horizontal line represents the expected amount of introgression, HI = 12.5 %, under neutrality. Vertical lines represent species-specific *D. flavomontana* (orange) and *D. montana* (blue) chromosomal inversions. Solid and dashed vertical lines show breakpoints of different inversions.

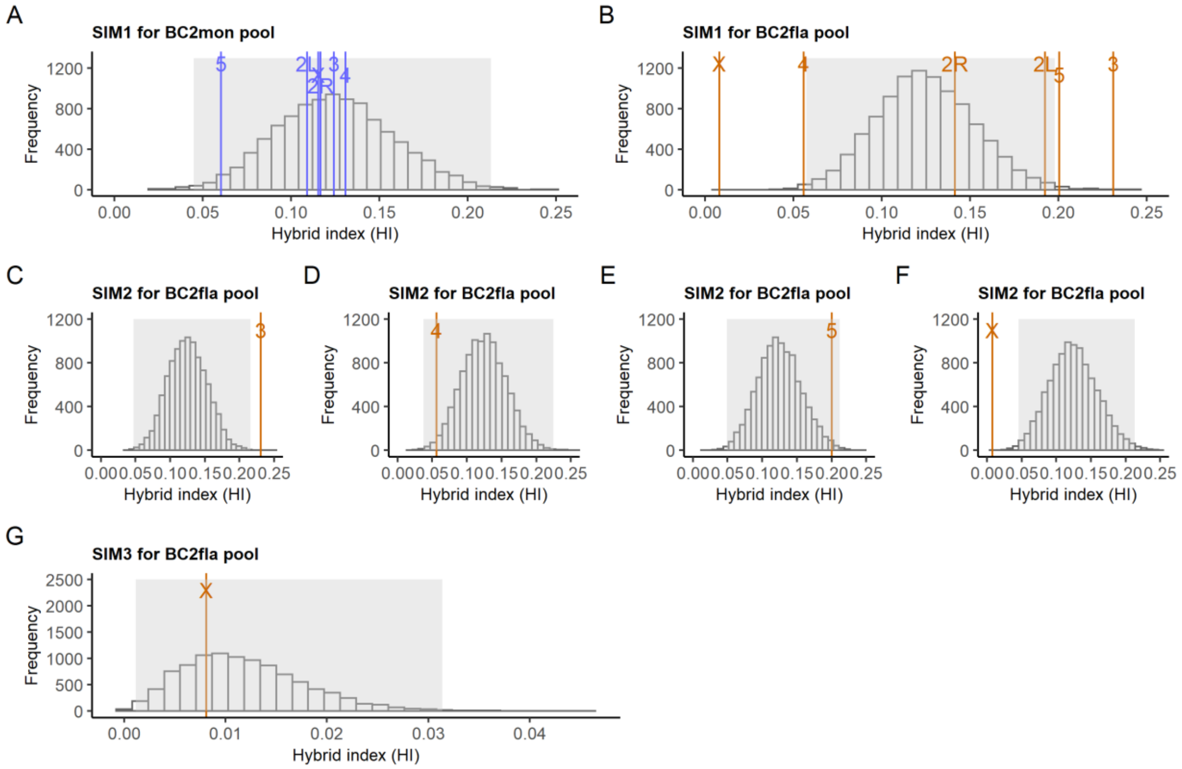


Figure S7. Hierarchical representation of the most meaningful simulations (10,000 replicates/simulation) of the 2nd generation backcross experiments towards *D. montana* (BC2mon) and *D. flavomontana* (BC2fla) (*D. montana* was used as a reference genome). The grey area of each figure represents Bonferroni corrected 5% and 95% quantiles and the space between them (regions beyond the area are statistically significant). Simulations under neutrality (SIM1) and the observed mean hybrid index (HI) of each chromosome for (A) $BC_2$mon pool and (B) $BC_2$fla pool. Simulations under neutral inversions (SIM2) and observed mean HI of $BC_2$fla pool for (C) the 3$^{rd}$ chromosome, (D) the 4$^{th}$ chromosome, (E) the 5$^{th}$ chromosome, and (F) the X chromosome. (G) Simulations involving inversions with a single locus against introgression (SIM3) and observed mean HI for the X chromosome of $BC_2$fla pool.

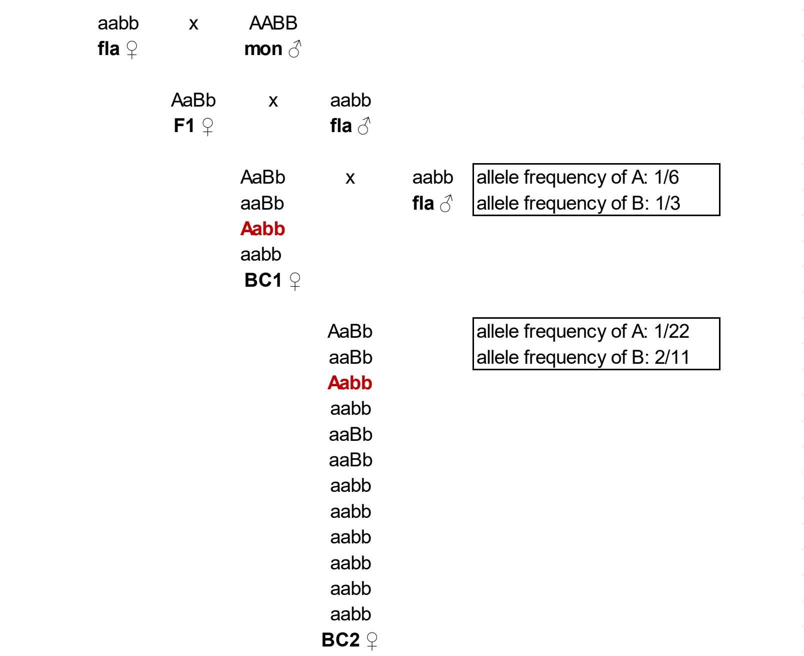


Figure S8. Illustration of the expected allele frequencies of *D. montana* (A and B) after selecting a pairwise dominant-recessive incompatibility in hybrids backcrossed to *D. flavomontana*. A/a locus is dominant and B/b locus is recessive (note that we investigated only SNPs that were differentially fixed between the species). Red genotypes, involving dominant *D. montana* locus interacting with recessive *D. flavomontana* locus, are either sterile or inviable and will not contribute to the allele frequencies.
